# Genomic, Proteomic and Phenotypic Heterogeneity in HeLa Cells across Laboratories: Implications for Reproducibility of Research Results

**DOI:** 10.1101/307421

**Authors:** Yansheng Liu, Yang Mi, Torsten Mueller, Saskia Kreibich, Evan G. Williams, Audrey Van Drogen, Christelle Borel, Pierre-Luc Germain, Max Frank, Isabell Bludau, Martin Mehnert, Michael Seifert, Mario Emmenlauer, Isabel Sorg, Fedor Bezrukov, Frederique Sloan Bena, Hu Zhou, Christoph Dehio, Giuseppe Testa, Julio Saez-Rodriguez, Stylianos E. Antonarakis, Wolf-Dietrich Hardt, Ruedi Aebersold

**Affiliations:** Department of Pharmacology, Yale Cancer Biology Institute, Yale University School of Medicine, West Haven, CT 06516, U.S.A; Joint Research Center for Computational Biomedicine (JRC-COMBINE), Faculty of Medicine, RWTH Aachen University, Aachen, Germany; Department of Biology, Institute of Molecular Systems Biology, ETH Zurich, Zurich, Switzerland; Institute of Microbiology, ETH Zurich, 8093 Zurich, Switzerland; Department of Genetic Medicine and Development, University of Geneva Medical School, and University Hospitals of Geneva, 1211 Geneva, Switzerland; Department of Experimental Oncology, European Institute of Oncology, Milan, Italy; Institute for Medical Informatics and Biometry, Carl Gustav Carus Faculty of Medicine, Technische Universität Dresden, Dresden, Germany; National Center for TumorDiseases, Dresden, Germany; Biozentrum, University of Basel, Klingelbergstrasse 50/70, 4056 Basel, Switzerland; Service of Genetic Medicine, University Hospitals of Geneva, Geneva, Switzerland; Department of Analytical Chemistry and CAS Key Laboratory of Receptor Research, Shanghai Institute of Materia Medica, Chinese Academy of Sciences, China; Department of Oncology and Hemato-Oncology, University of Milan, Italy; European Molecular Biology Laboratory - European Bioinformatics Institute, Wellcome Genome Campus, Cambridge, United Kingdom; iGE3 Institute of Genetics and Genomics of Geneva, Switzerland; Faculty of Science, University of Zurich, 8057 Zurich, Switzerland

## Abstract

The independent reproduction of research results is a cornerstone of experimental research, yet it is beset by numerous challenges, including the quality and veracity of reagents and materials. Much of life science research depends on life materials, including human tissue culture cells. In this study we aimed at determining the degree of variability in the molecular makeup and the ensuing phenotypic consequences in commonly used human tissue culture cells. We collected 14 stock HeLa aliquots from 13 different laboratories across the globe, cultured them in uniform conditions and profiled the genome-wide copy numbers, mRNAs, proteins and protein turnover rates via genomic techniques and SWATH mass spectrometry, respectively. We also phenotyped each cell line with respect to the ability of transfected *Let7* mimics to modulate *Salmonella* infection.

We discovered significant heterogeneity between HeLa variants, especially between lines of the CCL2 and Kyoto variety. We also observed progressive divergence within a specific cell line over 50 successive passages. From the aggregate multi-omic datasets we quantified the response of the cells to genomic variability across the transcriptome and proteome. We discovered organelle-specific proteome remodeling and buffering of protein abundance by protein complex stoichiometry, mediated by the adaptation of protein turnover rates. By associating quantitative proteotype and phenotype measurements we identified protein patterns that explained the varying response of the different cell lines to *Salmonella* infection.

Altogether the results indicate a striking degree of genomic variability, the rapid evolution of genomic variability in culture and its complex translation into distinctive expressed molecular and phenotypic patterns. The results have broad implications for the interpretation and reproducibility of research results obtained from HeLa cells and provide important basis for a general discussion of the value and requirements for communicating research results obtained from human tissue culture cells.

## Introduction

The independent verification of research results is the bedrock of experimental research. In 2011–2012 scientists at pharmaceutical companies reported that they were unable to replicate 80-90% of the findings in landmark research papers^1,2^, thus documenting a “reproducibility crisis” in the life sciences. A survey conducted by *Nature* in 2016 with 1,576 researchers suggested that 52% of participants agree on the existence of a significant “crisis” of reproducibility, and that publications in biology and medicine were deemed less reproducible than those in chemistry and physics^3^. In the last five years the scientific community has started to address the problem by promoting transparency in scientific publications^4^, analyzing and identifying key factors that influence reproducibility^5–8^, and by independently replicating selected results from high-profile papers, e.g. in the field of cancer biology^9–11^. These efforts have already resulted in better documentation of protocols and reagents in the scientific literature and, through mandating the deposition of primary data in public repositories, in an improved ability to independently verify conclusions derived from published data. Whereas these initial steps are useful, they are clearly not sufficient. Multiple factors can contribute to poor reproducibility, including poor study design, improperly applied statistical methods, lack of data sharing and transparency, the incompetence of researcher, inappropriate or poorly reproducible measurement methods, incomplete description of experimental details and incorrect or mislabeled reagents^7,8^. Apart from these technical issues, the complexity inherent in biological systems further challenges the reproducibility of life science research. At present it is generally unknown how genomic and environmental perturbations affect the molecular makeup and ensuing phenotypic response of a cell or organism, and how (genetically) different cells or organisms react to identical perturbations.

Human cancer cell lines are widely used in biological and biomedical research. They represent a source of biological material that can be perpetually regenerated, is technically straightforward and economical to maintain and is assumed to deliver reproducible results. For these reasons human cancer cell lines are frequently preferred over primary cultures or *in vivo* research. Several recent studies have highlighted significant problems with the use of human cancer cell lines, including cell line misidentification, cross-contamination and poor annotation, which altogether impair the reproducibility of results obtained from such cell lines between labs^12–15^. Consequently, the correct nomenclature of cultured cells has received significant attention. For instance, short tandem repeat profiling has been proposed to correctly identify cells used^15, 16^. However significant, experimental artifacts like contamination or mislabeling are only a part of the problem and could, in principle, be alleviated by careful experimentation. To date it has not been thoroughly explored to what extent cultured cell lines remain genetically stable over time and to what extent potential genotypic variability induces proteotype and phenotype variations in a given cell line with the *“same name*” across laboratories. This question deserves critical consideration because anecdotal, non-systematic observations suggest that many cultured cell lines might be genomically unstable^17, 18^. Therefore, even careful experimentation cannot fully alleviate the problem.

In this study we have quantified the degree of genomic variability, its effects on multiple levels of gene expression and a phenotypic response using HeLa cells as a model. HeLa cells originated from a cervical cancer tumor in a patient—Henrietta Lacks—in 1951^19^ and have been extensively used since. Several reasons led us to choose HeLa cells for this study. *First,* HeLa is the first successfully immortalized human cell line which has widely impacted biological studies. These include several studies for which Nobel Prizes were awarded, such as the development of the polio vaccine^20^, the discovery of telomeric activity^21^, and the linkage between human papilloma virus and cervical cancer^22^. More than 97,000 publications—nearly 0.3% of all publications curated in PubMed—have directly used or directly referenced HeLa cells. *Second,* HeLa cells have been reported for decades to contaminate and dominate other cell lines^23, 24^ and conversely, they are less likely to be contaminated themselves, minimizing the possibility of cell misidentification for HeLa-focused studies^15, 16^. *Third*, due to extensive genome instability during passaging and between labs HeLa cells were reported to contain an exceptional number of genomic variants^18,25–28^. For example, the currently widely used HeLa variants include HeLa CCL2, the “original” HeLa cell line; HeLa S3 (also called CCL2.2), the third clone isolated from an early HeLa culture; and HeLa Kyoto, the popular Kyoto version. Whole genome sequencing of HeLa Kyoto^29^ and HeLa CCL2^30^ strains were carried out separately, and significant variation of sequence, gene copy numbers, and chromosomes were reported between the two^30^. Furthermore, different HeLa Kyoto clones between labs were reported to vary in terms of mRNA expression^18^. However, little is known about the manner by which such genomic variability affects the proteomes or phenotypes of the different HeLa strains between individual labs, the effect of successive passages within a stock-derived line on its molecular make up, and how these would affect a biological research outcome^31, 32^.

Proteins catalyze and control most biological processes. The concentration and the synthesis and degradation rates of proteins in many scenarios cannot be predicted from corresponding mRNA abundance^33–35^. Over the last decade the proteome has become precisely measurable and it has been recognized that the acute state of the proteome—the *proteotype*—is a useful indicator of the cellular state and thus important for the understanding of disease phenotype, development, pharmacological responses, and many other aspects of life science research^36, 37^. The last five years have also witnessed significant technological improvements in mass spectrometry (MS)-based proteomics regarding proteome coverage and measurement reproducibility of large sample cohorts. In particular, data-independent acquisition (DIA) methods, exemplified by SWATH mass spectrometry (SWATH-MS) combined with targeted data extraction has achieved unprecedented reproducibility, generating quantitative matrices for thousands of proteins measured across multiple samples^37–39^. Recently, the statistical strategies controlling the quality of protein identification and quantification in large-scale

SWATH-MS have matured^40, 41^, and SWATH-MS has proven to be reproducible in a cross-lab study for ~5,000 proteins quantified in mammalian cells^42^. We therefore chose the SWATH-MS method to quantify the steady state proteome of 14 HeLa cell line variants collected from 13 labs. We related the quantitative protein profiles to copy number profiles, transcript profiles, protein turnover rates^43–45^ and a phenotypic response. We posit that the results obtained in this study highlight the biological complexities of commonly used human cancer cell lines and provide an important basis for a general discussion of the value and the communication requirements of ensuing results.

## Results

### HeLa cell variants from different laboratories have different and rapidly evolving genotypes

To survey the genomic, proteomic and transcriptomic similarity of HeLa cells across research groups, we reached out to 13 laboratories to provide samples of HeLa cells used in their group (**Figure 1**). The cell lines were numbered randomly from HeLa 1 to HeLa 13. Four laboratories shared the 7^th^ Passage (P7) of an initial HeLa CCL2 cell line culture (the original version of HeLa, directly purchased from ATCC). These P7 cells were then separately cultured in three labs until Passage 15 or 20 (P15 or P20), a stage where cells are generally regarded as still acceptable for biology research (HeLa 6, 7, and 13) (**Figure 1a**). The fourth group receiving the common P7 cells provided both P7 (HeLa 14) and P50 cells (HeLa 12) which allowed us to identify molecular alterations occurring within the timeframe of ca. 3 months. In total the 14 HeLa cell variants comprised seven HeLa CCL2 lines, one HeLa S3 line, six HeLa Kyoto lines including one line of unknown provenance - HeLa 3 - which was later identified as HeLa Kyoto using data from this study. To eliminate potential differences caused by varying culturing conditions and techniques between laboratories and to generate the number of cells (>10^7^cells per line) required for the multi layer molecular analyses, we centrally cultured all 14 cell lines for an additional three passages in our group. To minimize experimental variability, the same researcher cultured the cells using the same culture media and protocol.

**Figure 1.**
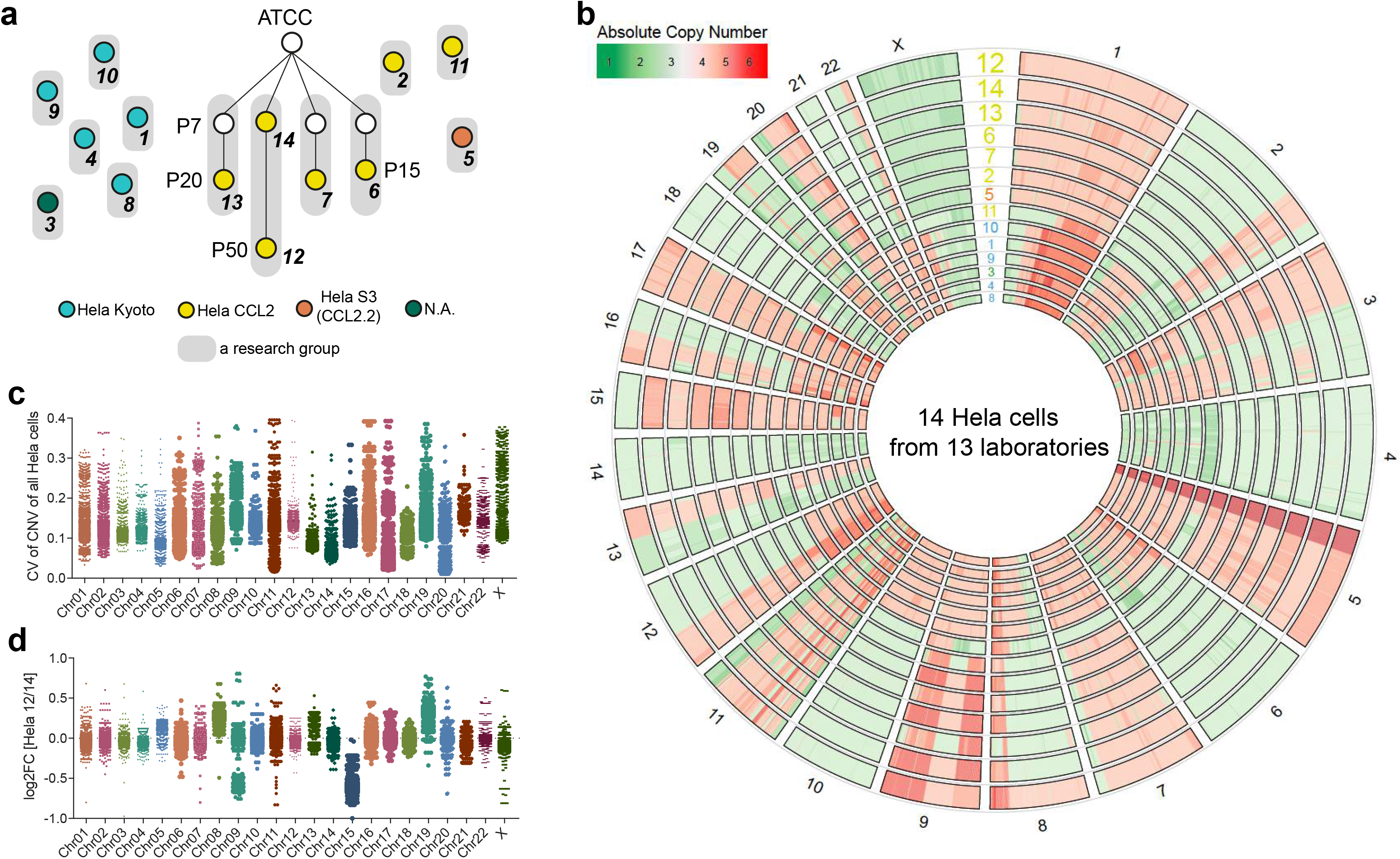
HeLa cell lines from different laboratories showed varied and evolving genotypes. **(a)** HeLa cell variants collected from 13 laboratories. ATCC, American Type Culture Collection organization; P, Passages. Px with x denotes the passage number of cultured cells. **(b)** Circus plot of raw absolute copy numbers across all 14 HeLa cells. **(c)** Coefficient of variation (CV) distribution of all genes encoded by different chromosomes among all cell lines. **(d)** Log-fold-changes of gene copy number changes between HeLa 12 and HeLa 14 were visualized at each chromosome.

As a starting point for the evaluation of HeLa cell heterogeneity, we measured gene copy number variation (CNV) by array-CGH (aCGH). Previous studies reported genome instability within and between different HeLa strains_18,25-28_. To investigate and display the ploidy variation of the strains studied we projected the CNV data onto the reference human genomic map (**Figure 1b**). We discovered pervasive CNV differences organized by domains, large chromosomal segments, and even whole chromosomes. Particularly notable were ploidy changes at Chromosomes (Chr) 1, 2, 6, 9, 10, 17, 19, 21, 22, and X. (**Figure 1b**). On average, the HeLa genome of all cells tested had an overall hypertriploid state, as reported^29, 30^. However, distinctive patterns differentiated the Kyoto and CCL2 lines: HeLa CCL2 lines had 1.87-times as many genes with two copies and 0.7-times as many genes with three copies compared to the Kyoto lines, suggesting that the original HeLa CCL2 cells have a generally smaller genome than Kyoto variants (**Supplementary Figure 1**). Even within the CCL2 and Kyoto groups, significant CNVs could be observed between samples provided from different laboratories, albeit at a smaller scale, exemplified by the distal locus on Chr8 (**Figure 1b**). Moreover, HeLa 11 showed a deviating pattern in multiple chromosomes compared to other HeLa CCL2 cells. The distribution of coefficient of variation of all the genes encoded by different chromosomes among all cell lines revealed genome-wide DNA dosage differences, e.g., Chr13, 14, and 18 seemed to be relatively stable among HeLa cell variants tested (**Figure 1c**).

To investigate the rate of genome variability progression, we investigated the CNV divergence between HeLa 14, the original ATCC HeLa CCL2 cell line at P7, and HeLa 12, the direct descendent of HeLa 14 at P50. The observed fold-changes were visualized at each chromosome (**Figure 1b**, outermost circles & **Figure 1d**). Somewhat surprisingly, after ~3 months of continuous passaging, the HeLa cells tested already gained or depleted entire chromosome copies or large chromosome domains. Examples are the copy gains of whole Chr8 and loss of Chr15, as well as a two-thirds gain of Chr 19 and a partial (30%) gain of Chr 9.

In summary, we observed, as a likely consequence of genomic instability, a considerable degree of large scale CNV across HeLa cells cultured in different labs, even among strains with the same annotation.

### Diversity of gene expression patterns at steady state

We then posited that these CNVs would impact gene expression patterns of the respective HeLa strains at steady state. Therefore, we further performed transcriptomic profiling by mRNA sequencing, steady state proteomic profiling by SWATH-MS, and protein turnover rate determination by pulse-chase SILAC (pSILAC) labeling followed by SWATH-MS. pSILAC measures the incorporation rate of isotopically heavy amino acids into newly synthesized proteins^43–45^. Experts in the respective techniques, blinded to the cell identities, carried out the measurements at each layer of gene expression, thus contributing to a unique multi-layer dataset of related cultured cells (**Figure 2**).

**Figure 2.**
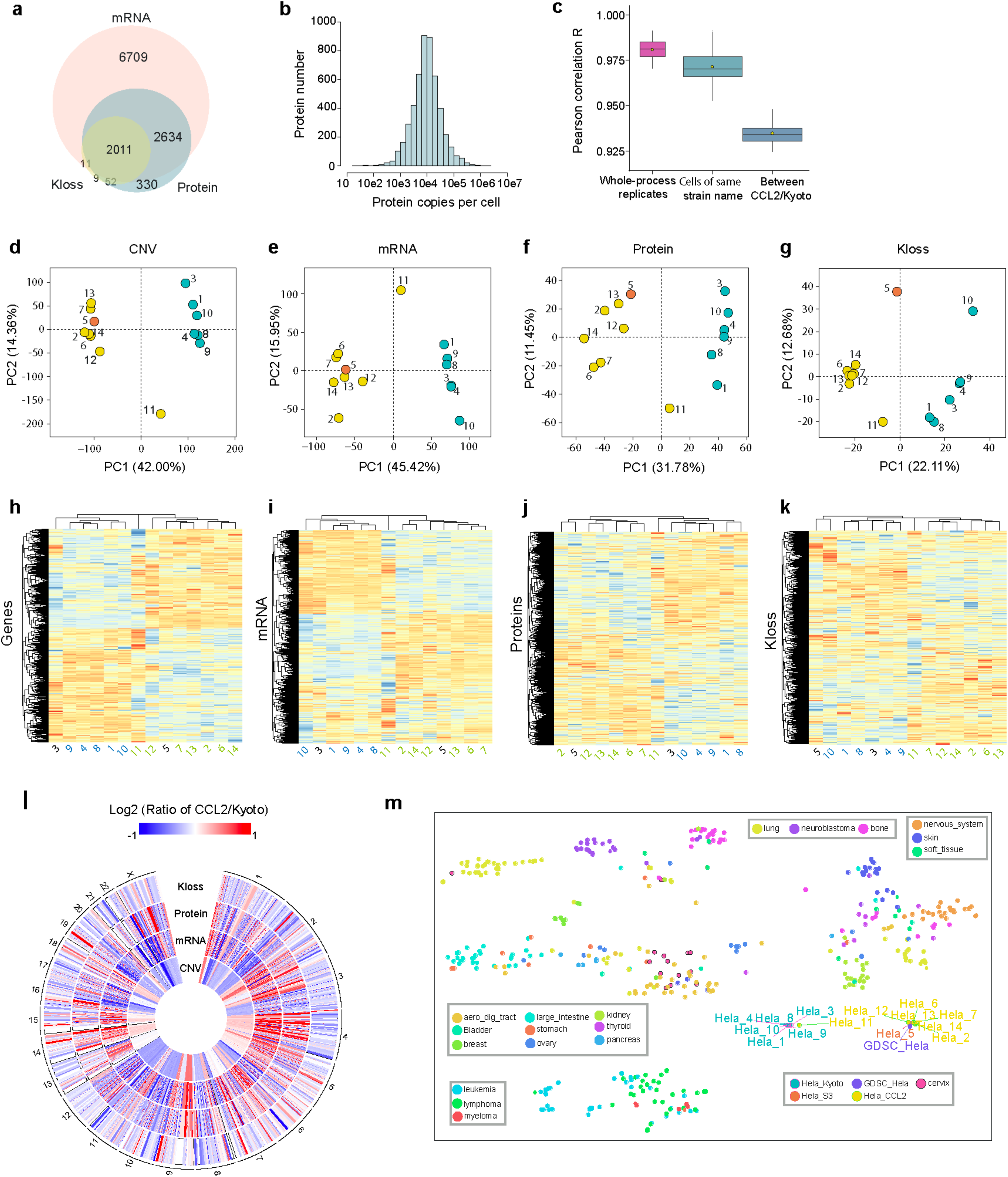
Heterogeneous transcriptome, proteome, and protein turnover profiles between HeLa cell lines across laboratories. **(a)** Expression of genes measured at each layer. **(b)** Label-free absolute quantification of averaged protein copies per HeLa cell. **(c)** Quantitative reproducibility of SWATH-MS is sufficient to distinguish whole process replicates and HeLa cells from different labs. **(d-g)** Principal component analysis (PCA) of gene-wide CNV, transcripts (N=11,365), proteins (N=5030), and Kloss (N=2084) of all HeLa cell lines. *Kloss,* protein loss rate, the proxy of protein turnover rate. **(h-k)** Hierarchical-clustering analysis (HCA) of the data. **(l)** HeLa CCL2/Kyoto ratios. Note the fold-change greater than 2 are labeled by extreme colors. The data is proteome centric, i.e., data matched to available proteomic identifications. **(m)** t-SNE analysis of our HeLa transcriptomic data combined with for 1,001 molecularly annotated human cancer cell lines from Iorio et al.

Collectively, we quantified transcripts for 11,365 genes (after filtering by average RPKM>1 among all 14 lines, **Figure 2a**). Using the state-of-the-art SWATH-MS measurements^41,46,47,48^, we confidently quantified 5,030 proteins across all samples (1% peptide and protein FDR controlled by the PyProphet tool^41^, for details see **Method & Supplementary Figure 2**). By absolute label free quantification, the abundance of the 5,030 proteins was determined to center at 10,000 copies per cell, spanning a range of abundance from 100 to >10^6^ copies per cell (**Figure 2b**). Using pSILAC SWATH-MS, we quantified the proxy turnover rate (i.e., *Kloss,* see Method) of a consistent set of 2,084 proteins in each HeLa cell line. Together, these analyses yielded 4,656 genes with both mRNA and protein measurements, and 2,011 genes with all three layers of data, presenting a substantial resources for gene expression studies in cells (**Figure 2a & Supplementary Figure 3**).

To assess the quantitative reproducibility of SWATH-MS, we performed correlation analysis (**Figure 2c**). The binary Pearson correlation coefficient R was on average 0.981 (95% CI, 0.9769–0. 9847) for whole process replicates (i.e., replicates of independently cultured cell batches of the same HeLa aliquot). This value is significantly higher (*P* < 0.0001, Mann-Whitney test) than the correlations observed between cells of the same HeLa group but from different laboratories (i.e., within CCL2 and Kyoto group), which showed an average coefficient R of 0.971 (95% CI, 0.9693–0. 9733). As expected, the comparisons between CCL2 and Kyoto cells showed the lowest binary correlations (R=0.935; 95% CI, 0.9332-0.9362). These comparisons suggest that SWATH-MS achieved a high and consistent reproducibility that is sufficient to distinguish whole process replicates and HeLa cells from different labs, even independent from the genomic data.

To compare HeLa cells at each omics layer, we performed simple quantile normalization for the CNV, mRNA, protein, and *Kloss* data, respectively, and visualized their variation through PCA (**Figure 2d-g**) and unsupervised hierarchical-clustering analysis (HCA, **Figure 2h-k**). In PCA, the first principle component of each of the CNV, mRNA, protein, and *Kloss* levels already separated HeLa CCL2 and HeLa S3 (HeLa 5) cells from HeLa Kyoto cells. In HCA, these two major groups also emerged as separated clusters at all levels tested. Our results thus suggest that, *first,* HeLa CCL2 and Kyoto variants substantially differ at every level of gene expression, *second,* that HeLa S3 (HeLa 5) cells are genomically closer to HeLa CCL2 than to Kyoto variants (**Figure 2h**) a pattern that is recapitulated at the mRNA, protein, and proteostasis level (**Figure 2j-l**) and *third*, compared to other CCL2 cells, HeLa 11, which has an unusually distinct CNV pattern, also showed distinct discrepancies of transcriptional and post-transcriptional gene expression. HeLa 11 resides between CCL2 and Kyoto groups in PCA, although it is closer to the CCL2 cluster in HCA at all levels. To confirm the cell identity of HeLa 11 (declared by the donating lab), we performed pairwise cell line concordance analysis^49^ by the deep mapping of single nucleotide variants (SNV) extracted from our RNA-seq dataset (**Supplementary Figure 4**). According to this analysis, all the HeLa cells including HeLa 11 have a very similar extent of similarity for SNV, suggesting they are *bona fide* HeLa cell lineages. To sum up, HeLa cells can be distinctive in gene expression patterns.

We next explored the observed difference between HeLa CCL2 and Kyoto groups in more detail. We firstly aligned the CCL2/Kyoto ratios at CNV, mRNA, protein, and *Kloss* levels with the respective loci in the human genome (**Figure 2l**). We found that the transcriptome and proteome differences largely followed the gene copy number imbalance reflected by CNV ratios, whereas the Kloss ratios showed a higher degree of variation. We then examined how differences between CCL2 and Kyoto compare to the differences between other human cell lines of different types and origins. We referred to a previous published data set which contains the RNA-seq data for 1,001 molecularly annotated human cancer cell lines (GDSC panel)^50, 51^, and used t-SNE analysis to plot the variance at the transcriptomic level between HeLa cells and the GDSC cell lines (**Figure 2m**). We found that based on transcriptomic data, HeLa CCL2 and Kyoto groups are as distinct from each other as are cancer cell lines from different tissue types. Furthermore, the published HeLa cell in the GDSC panel (the strain identity not mentioned) clustered with our CCL2 cells, indicating that it is likely a HeLa CCL2 cell line. To corroborate this analysis, we performed t-SNE for proteome and Kloss data by comparing HeLa cells to the primary skin fibroblast cells isolated from 11 Down Syndrome and 11 control individuals and a monozygotic twin pair discordant for trisomy 21 for which the same proteomic techniques (i.e., SWATH-MS and pSILAC-SWATH) were employed^45^ (**Supplementary Figure 5**). We found that all HeLa cell variants were clearly separated from fibroblast cells by both protein abundance and turnover rates. Interestingly, the quantitative proteomic variance between HeLa cells which is associated with substantial karyotypic differences (**Figure 1**) is larger than that caused by an additional chromosome 21, the smallest human autosome, but appears smaller than the variance between genetically unrelated individuals.

Taken together, HeLa cells from different labs showed substantially diverse gene expression patterns, and the quantitative variation observed between HeLa cells is comparable to the variability observed between cells of different tissue origin.

### Gene expression patterns evolve with cell passages

HeLa cells like most cell lines commonly used in research are immortalized. Although the best practice in a biological lab is to restore cell aliquots from the earliest generation as frequently as possible and to thaw them upon use, it is practically difficult to know the precise history of the cells being used in an experiment, e.g., how many generations or passages the current cells have already been through since being initially obtained from ATTC. Also, it is not clear how much functional variation will be introduced during passaging of cells with unstable genome, as is the case for HeLa^18, 25,28^. We therefore assessed the extent of gene expression difference between the HeLa 12 and 14 cells, representing P7 and P50, respectively, of the same starter culture. We compared the mRNA-seq data acquired from three biological replicates each of HeLa 12 and 14. Strikingly, 731 transcripts (~6.4% of the total transcripts confidently profiled) showed significantly altered expression between HeLa 14 and 12 (adjusted P<0.01 by edgeR^52^, fold-change>2) (**Figure 3a**). Among these, 415 transcripts showed increased expression levels, whereas 316 transcripts showed decreased expression levels in P50 cells. These 731 transcripts were significantly enriched in several distinct biological processes annotated in GO (**Figure 3b**). These include system development (GO:0048731, adjusted P=2.3^-21^), response to lipid (GO:0033993, adjusted P=2.5^-20^), positive regulation of biological process (GO: 0048518, adjusted P=1.6^-15^), and cell surface receptor signaling pathway (GO:0007166, adjusted P=4.2^-15^). These results argue that investigations focusing on these functions or processes might potentially lead to inconsistent results if HeLa cells like HeLa 12 and HeLa 14 are used within or between labs.

**Figure 3.**
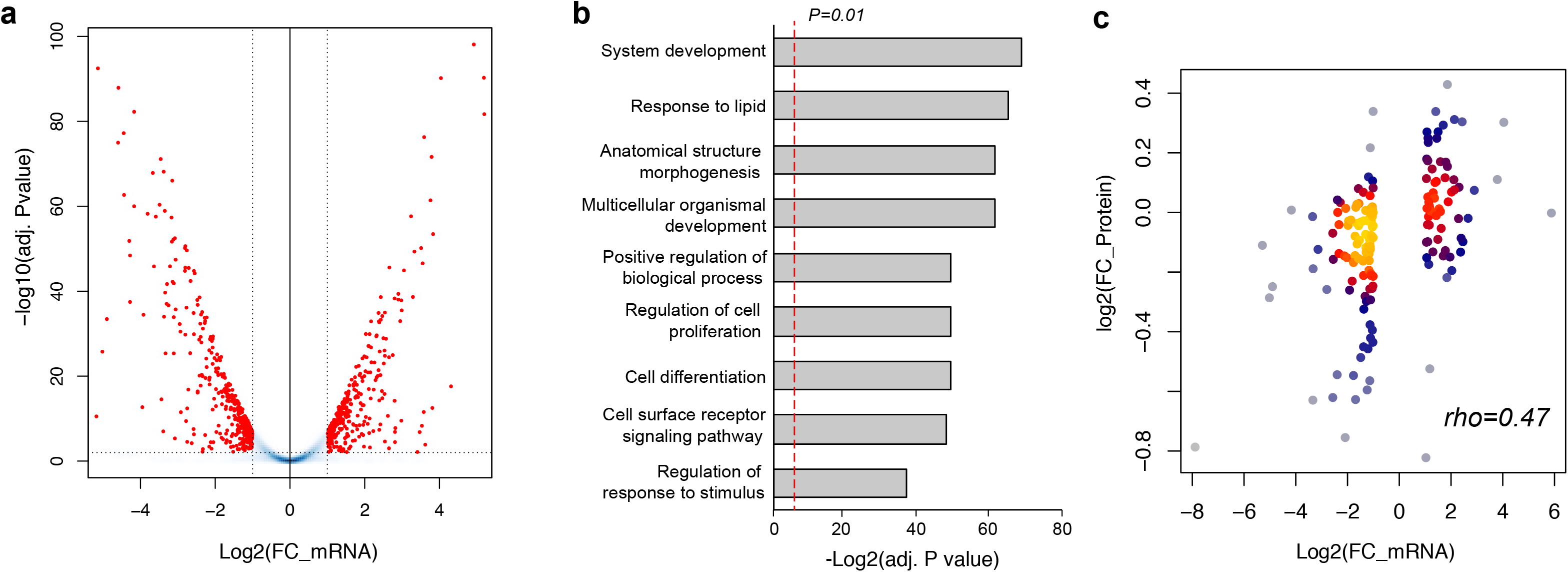
Gene expression comparison between HeLa 12 and 14 representing 3-month passaging. **(a)** Volcano plot of 3 vs. 3 biological replicates by transcriptomics data. Note the red circles denote 415 transcripts showed increased expression levels, whereas 316 transcripts showed decreased expression levels in HeLa 12 (P50) compared to HeLa 14 (P7). **(b)** Representative GO biological processes enriched by differential transcripts. **(c)** Spearman correlation analysis between quantitative data of mRNA and protein fold-changes revealed the translated effect of differential mRNA abundances. Spearman’s *rho* is shown.

We next asked whether these differential transcripts profiles were translated into differential protein profiles. Using SWATH-MS, we were able to quantify proteins corresponding to 166 transcripts out of 731 differential transcripts. The mRNA~protein quantitative correlation between HeLa 12 and 14 was significant (Spearman’s rho=0.47, **Figure 3c**), suggesting that the differential transcripts detected by comparing P7 and P50 significantly impact the quantitative proteome and thus, diverse cellular functions (**Supplementary Figure 6**).

### Global processes shape HeLa proteotypes

The multi-layer omics data set obtained from the 14 HeLa cell lines allowed us to systematically analyze global proteome expression control mechanisms including gene dosage compensation^34, 45,53–56^. In addition, we could ask how the cells reconcile the dosage effect of large-scale CNVs along chromosomes with protein requirements and constraints in essential metabolic pathways.

**Figures 4a-c** visualize the quantitative differences of CNV and mRNA and protein levels between HeLa cells after normalizing the data to the mean of the respective values. Chromosome 1 and 11 are shown as representative examples. It is apparent that the numerous CNVs led to significant mRNA changes that did not translate into the corresponding protein levels. Also, there was a decrease in total variance summarized at each layer from mRNA to protein and to protein turnover rate (**Supplementary Figure 7**). These results suggest the existence of global buffering mechanisms along the axis of gene expression from transcripts to proteins^34^

**Figure 4.**
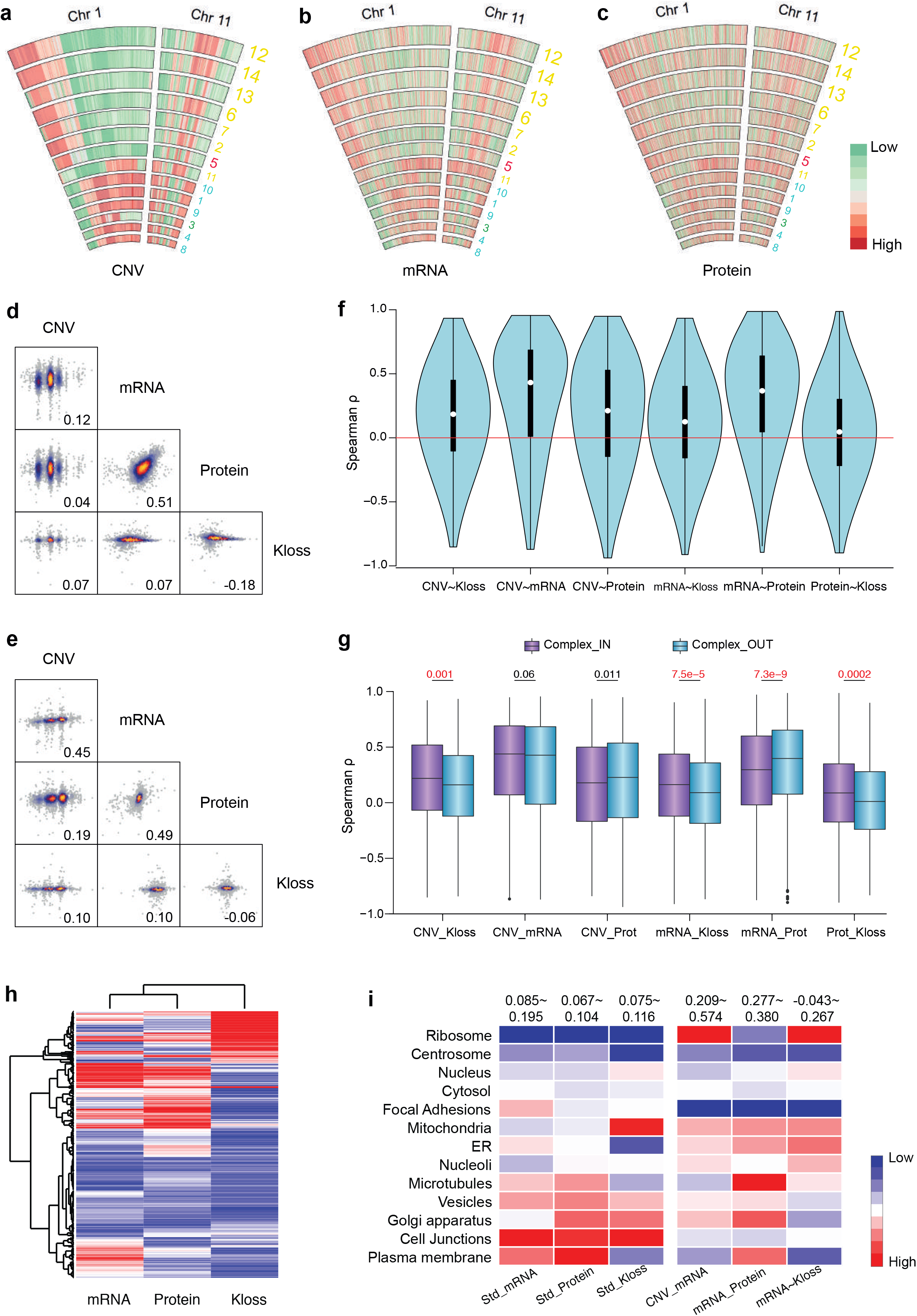
Global processes affecting HeLa proteotypes. **(a-c)** Relative differences for CNV, mRNA, and protein levels between HeLa cells after data centralization. Chromosome 1 and 11 are shown as examples. **(d)** Across-gene correlation between CNV, mRNA, protein, and *Kloss* values using absolute-scale data HeLa 1 (a HeLa Kyoto cell). **(e)** Within-gene correlation between layers using the relative quantification data between HeLa 1 and HeLa 14, the ATCC CCL2. **(f)** Gene specific, cross-cell line Spearman rho between layers. **(g)** Cross-layer correlations between those proteins annotated in any stable complex through CORUM database (“complex_IN” group) and those are not (“complex_OUT” group). **(h)** Gene expression variability calculated by standard deviation from the average at each level suggests the regulation preferences. **(i)** Organelle perspective of gene expression variability and correlation.

To dissect post-transcriptional protein expression control, we and others have emphasized the importance of analyzing across-gene, within-gene, and gene specific correlations between omics data sets^34, 35^. Thus we performed correlation analyses between CNV, mRNA, protein, and *Kloss* values comparing both, across-gene and within-gene patterns. As an example, the absolute-scale data from an arbitrary reference cell e.g. HeLa 1 (a HeLa Kyoto cell) were correlated between omics layers in the across-gene manner. (**Figure 4d**). HeLa 1 was further compared relatively to other lines e.g. HeLa 14, a HeLa CCL2 line by correlating the fold-changes at each layer (**Figure 4e**). The mRNA~protein correlation was quantitatively strong for both absolute and relative scales (Spearman correlation coefficient rho=0.51 and 0.49), reinforcing the previous notion that protein levels at steady state are primarily determined by mRNA levels^34, 57^. Furthermore, the protein turnover rate, *Kloss,* was found to fine-tune gene expression in general, showing small but non-negligible positive correlations to CNV and mRNA levels. The absolute protein~Kloss correlation *rho* was −0.18 (**Figure 4d**), denoting the fact that highly abundant proteins are less strongly regulated by protein degradation rates than proteins expressed at lower levels^58^. Notably, CNV heavily determines the mRNA levels when we consider the relative difference between two cell lines (rho=0.45), but is only marginally significant for predicting mRNA copies within one cell line (*rho* =0.12). Similar trends were found for protein levels *(rho* =0.19 vs. 0.04). This illustrates the importance of considering both, across-gene and within-gene analyses.

To further evaluate gene-specific predictions we plotted the distribution of genome-wide Spearman’s *rho* calculated across the 14 HeLa cells. Consistently, we found the highest correlations between CNV~mRNA (averaged rho=0.27) and mRNA~proteins (averaged rho=0.31), suggesting the conserved information flow along the central dogma axis for most genes. The observed CNV~protein correlation was less strong (averaged rho=0.17) in part due to the interspersed mRNA level control. Furthermore, the CNV~Kloss and mRNA~Kloss correlations were 0.16 and 0.11, respectively, again supporting the notion that for many genes protein turnover functions as a buffering step, shaping the quantitative proteome of the HeLa cells tested (**Figure 4e-f**).

The maintenance of protein complex stoichiometry was reported as an efficient buffering mechanism against aneuploidy stress in different systems^34, 53, 54, 59^. We previously generated direct protein degradation measurements by pSILAC-SWATH to illustrate this proteostasis mechanism in human trisomy 21^45^. Here we tested the significance of protein complex stoichiometry control among 14 HeLa strains, by comparing the cross-layer correlations between those proteins annotated in any stable complex in the CORUM database (“complex_in” group) and those that are not annotated as complexes in CORUM (“complex_out” group) (**Figure 4g**). We found that the CNV~mRNA correlations are not significantly different between “complex_in” and “complex_out” groups (P=0.064, Mann-Whitney test). In stark contrast, the “complex_in” proteins tend to have significantly weaker mRNA~protein correlations (P=7.3^-9^) and much stronger mRNA~Kloss correlations (P=7.5^-5^) than “complex_out” proteins, demonstrating the post-translational constraint of protein complex stoichiometry control for “complex_in” proteins, a process that affected mRNA~protein correlation. All the distributions of CNV~Kloss, CNV~protein, and protein~Kloss correlations can be explained, in part by protein degradation mediated buffering of complex components (**Figure 4g**). We anticipate that the statistical significance of above buffering mechanisms will further increase if the protein complex components were derived directly from cell specific experiments e.g., by SEC fractionation^60^.

We next analyzed the layered gene expression variability of specific gene sets in the tested HeLa cells. HCA indicates that the variations at mRNA, protein and protein turnover levels do not affect genes equally (**Figure 4h**). The majority (~75%) of genes showed consistent variation extent between mRNA and protein levels, whereas *Kloss* particularly strongly affected a subset of genes. To determine whether such regulatory patterns are functionally relevant, we used a recently established subcellular atlas of the human proteome to annotate genes based on the organelle locations and cellular compartments^61^ (**Figure 4i**). Moreover, we distributed the between-layer correlations of genes, with CNV~mRNA indicating transcriptional regulation, mRNA~proteins indicating post-transcriptional regulation, and mRNA~Kloss indicating buffering against mRNA variation through protein turnover. This integrative plot (**Figure 4i**) provides biological insights. For example, the mRNAs and proteins that showed the highest and lowest degree of variation are associated to cell junctions and ribosomes, respectively. Further, the mRNA~protein correlation was at the low end for ribosomal proteins, demonstrating tight protein level control for this functional group and the important role of ribosomes in proteome remodeling. The highest mRNA~Kloss correlation for ribosomal genes suggests that the remodeling can be at least partially explained by protein turnover. Indeed, about 40% of cytosolic ribosome flipped the relative expression protein pattern between HeLa CCL2 and Kyoto cells, compared to CNV difference (**Supplementary Figure 8**). Moreover, both *Kloss* variation and mRNA~Kloss correlation of the mitochondrial proteins are among the highest observed. Thus, the protein turnover of the mitochondrial proteome is likely to be important in buffering aneuploidy stress and the resultant mRNA changes in and between HeLa cells, which agrees with our previous report on human trisomy 21^45^. Conversely, there was a very weak correlation of mRNA~Kloss for plasma membrane proteins, suggesting a weaker role of protein turnover in shaping variable membrane proteome between HeLa cells. Therefore, the extent of variation at the transcriptional, protein, and post-translational level seems to be frequently protein specific and determined by the protein functions and their cellular locations.

In summary, these results indicate that the translation of genotypic variability observed between HeLa cell variants into proteotypic variability is controlled by a multitude of global processes, including the control of protein complex stoichiometry and organelle specific protein turnover rates.

### HeLa proteotypic variability tightly links to phenotypic variability

We then analyzed phenotypes to better understand the consequences of the molecular heterogeneity of the tested HeLa cells. We firstly used molecular imaging methods, specifically DAPI staining to highlight the nucleus and RFP labeling for F-actin visualization. We also counted cell numbers to determine cell-doubling time for each centrally cultured strain. PCA of all extracted image features showed a clear separation between CCL2 and Kyoto cells, indicating that the cells are morphologically different in many aspects, such as the texture contrast of their actin structures (**Supplementary Figure 9**). Cell doubling times—derived from triplicate measurements—were also strikingly different between HeLa strains with extremes being 17.5 and 32.3 hours under identical culture conditions (**Figure 5a**). Generally, HeLa Kyoto cells grew faster than CCL2 cells (averaged doubling time, 21.1 vs. 28.2 hours; P=0.018, Welch’s t-test), an observation that has not been documented previously. Intriguingly, 63 proteins annotated by KEGG as related to cell cycle could be used to distinctively classify HeLa Kyoto, CCL2 or S3 lines, respectively (**Figure 5b**). We found that the absolute protein copies of Cyclin-dependent kinases 1, 2, 7 (CDK1, CDK2, and CDK7) were on average 57.7%, 46.8% and 81.2% higher in the Kyoto group than in the CCL2 group of cells (**Supplementary Figure 10**), possibly accelerating the cell doubling observed^62^, thus establishing a proteotype-phenotype link.

**Figure 5.**
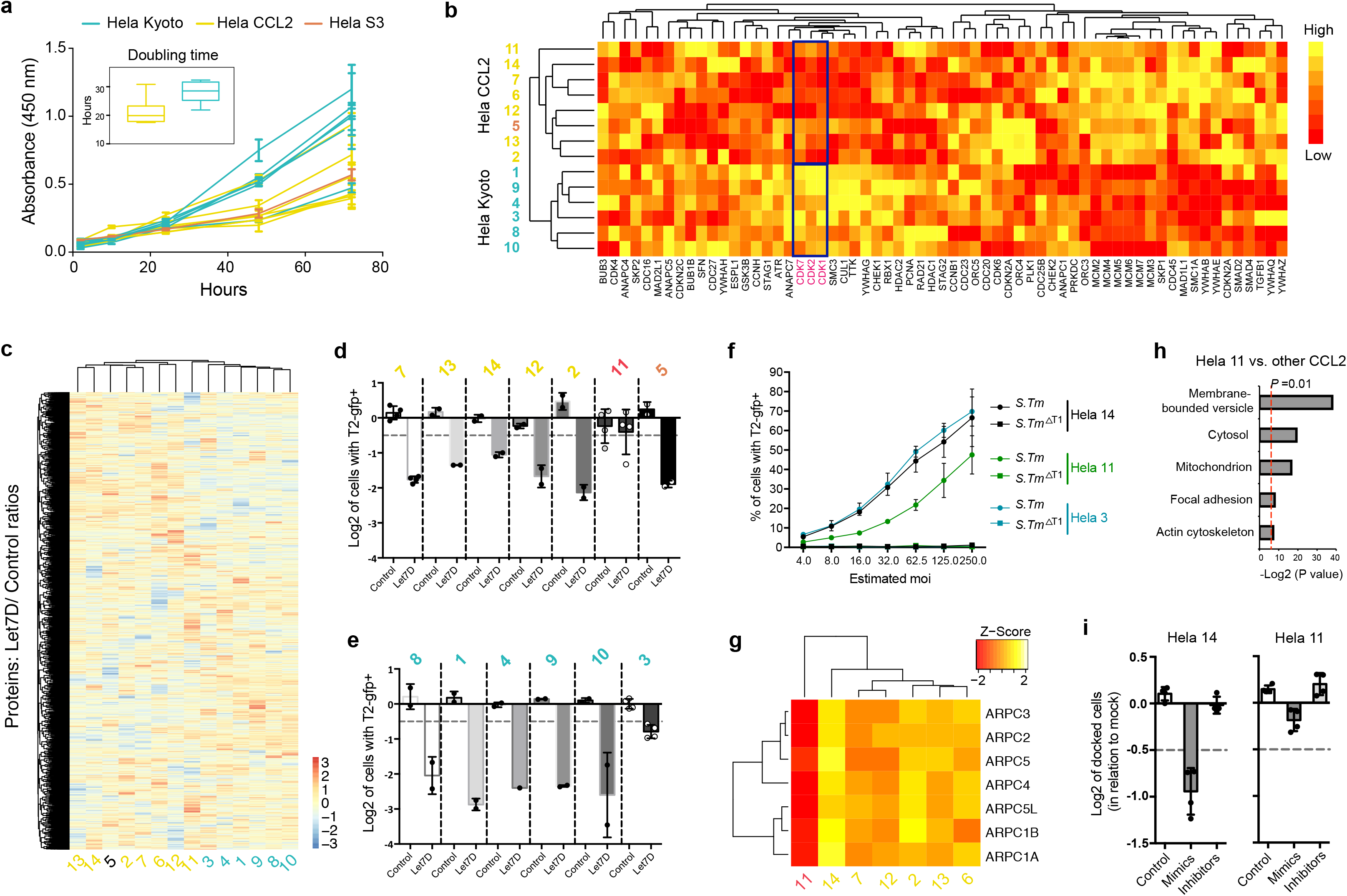
Proteotypes of HeLa cells tightly links to phenotypes. **(a)** Cell doubling time measurement suggests HeLa Kyoto group has faster cell doubling than HeLa CCL2 group. **(b)** 63 proteins annotated by KEGG to be involved in Cell Cycle process clearly differentiate HeLa CCL2 and Kyoto group, with all CDKs upregulation in Kyoto group, including the significant upregulations of CDK 1, 2, 7 highlighted. **(c)** HCA of the quantified all 5,030 proteins and their ratios between *Let7* treated vs. controls for each cell line. **(d)** *Salmonella Typhimurium* (S.Tm) infection rates quantified in HeLa CCL2 cell lines after *Let7 mimics* transfection which supposes to decrease the infection rate. See Supplementary Figure 11 for more information. **(e)** The infection rates in HeLa Kyoto cells. **(f)** HeLa 11 showed overall lower infection rates at the respective moi’s than HeLa 14 and HeLa 2. **(g)** HeLa 11 has the lowest expression of Arp2/3 complex among all HeLa CCL2 cells. **(h)** GO processes enrichment analysis for the 178 proteins that were differentially expressed between HeLa 11 and other CCL2 cells. **(i)** A less pronounced docking phenotype in HeLa 11 compared to HeLa 14 was confirmed.

Besides readily apparent phenotypes such as cell morphology and doubling time, we sought to inspect the consistency of the HeLa cells tested in their ability to respond to the same stimulus or perturbation. MicroRNAs (miRNAs) are known to suppress target gene expression in a sequence specific manner^63^. We therefore asked whether different HeLa cells could render different proteomic responses to perturbation via miRNA transfection. We selected mimics of *Let7,* which is a well-known, highly conserved miRNA that has been shown to play a central role in development and tumor suppression^64^”^66^. Using identical protocols, the set of 14 HeLa cell lines were transfected with *Let7* mimics or negative controls, respectively, and incubated for 72 hours. Using SWATH-MS we quantified 5,030 proteins and their abundance ratio between *Let7* treated vs. controls for each cell line. As for the other phenotypic measurements described above, the proteome-wide abundance ratios also clustered HeLa CCL2 and Kyoto cells into two respective groups. The only exception was HeLa 11 which was clustered between groups (**Figure 5c**). We specifically checked the protein ratio of Let7/control for those genes that were confirmed or predicted to be *Let7* targets (**Supplementary Figure 11**). SWATH-MS detected 108 of the top 500 most likely *Let7* gene targets according to TargetScan^67^ In the HeLa cells transfected with *Let7,* ~70% of the targets showed a decrease in abundance, indicating that *Let7* treatment was effective. Again, HeLa CCL2 and Kyoto versions were separated based on the abundance ratios of these 108 proteins, with 10 protein targets showing remarkably strain-dependent regulation upon *Let7* transfection in CCL2 and Kyoto. These results indicate that the HeLa cell lines tested, in addition to showing considerably different molecular gene expression patterns at steady state, also showed a different response to a specific perturbation. It is therefore likely that the different HeLa cell lines would also lead to different biological conclusions from experiments intended to identify *Let7* targets.

It was reported previously that the expression of *Let7* microRNAs is downregulated upon *Salmonella* infection in HeLa cells, a result that is indicative of a role of *Let7* in mediating the eukaryotic host response^65^. Using HeLa CCL2 (from ATCC; HeLa 14) we have identified four miRNAs including *Let7* for which mimics interfere negatively with *Salmonella Typhimurium* (*S.*Tm) infection and inhibitors interfere positively (**Supplementary Figure 12**). Further microbiological experiments suggested that the first docking step which precedes ruffling, entry and intracellular growth of *S.* Tm^68, 69^ was impaired in *Let7* treated HeLa CCL2 cells (**Supplementary Figure 12**). We therefore repeated the S.Tm infection experiment for all the HeLa cells transfected with *Let7d* (**Figure 5d-e**, only HeLa 6 was not included, see Methods). We found that the S.Tm infection rate was generally significantly reduced upon *Let7* transfection in most HeLa cell lines: the same concentration of *Let7* produced similar protective effect in most HeLa Kyoto cell lines. In contrast, the CCL2 group of cell lines showed a higher degree of phenotypic variability. HeLa 11 again showed an exceptional pattern in which the infection rate was not modulated by the *Let7* treatment (**Figure 5d**). To further explore this effect, we compared wild-type *Salmonella* and a non-invasive mutant (S.Tm ΔT1) for their ability to infect HeLa 11, HeLa 3, and HeLa 14 over a range of multiplicities of infection (moi) from 4 to 250. We found that HeLa 11 indeed showed lower overall infection rates at the respective moi’s compared to the other cells (**Figure 5f**). We therefore revisited the steady state proteome of HeLa 11. Reassuringly, we found that compared to other HeLa CCL2 cells, HeLa 11 has the lowest expression of the Arp2/3 complex which has a well-established role in bacterial internalization into host cells by initiating membrane ruffles^68–70^. Remarkably, all the seven subunits of the Arp2/3 complex that were quantified in the proteomic dataset followed the same pattern (**Figure 5g**), which could explain the reduced *S.Tm* infection in HeLa 11. We used the STRING database to perform gene set enrichment for the 178 proteins that were differentially expressed between HeLa 11 and other CCL2 cells (linear modeled test, adjusted *P* <0.05), and again we found the membrane-bounded vesicle (GO:0031988)^71^ was most significantly enriched (adjusted P=3.61E-12, **Figure 5h**). We finally performed a docking experiment^69^ by counting the S.Tm bacteria that remained bound to the respective HeLa cells after extensive washing. We confirmed a less pronounced *S.Tm* docking phenotype in HeLa 11 compared to HeLa 14, which may further indicate that HeLa 11 has a different membrane topology or composition (**Figure 5i**). Such membrane topology effects are known to affect the rate of S.Tm docking to host cells^72^. These results indicate that the different molecular response to *Let7* of the tested HeLa cells directly affected a bacterial infection phenotype.

To summarize, our data show that the set of proteins that constitute the cell line specific expression response provide useful information about the molecular basis underlying the observed phenotypic differences such as doubling time and miRNA modulated bacterial infection.

### A website for navigating HeLa cell line heterogeneity

Due to the good technical reproducibility and the matched CNV, mRNA, protein, and protein turnover data across varying HeLa samples, we consider the overall data set a high-quality resource for future HeLa related studies. We therefore generated a HeLa Proteome Website (https://saezlab.shinyapps.io/HeLa_App/) to navigate and contemplate the data basis of the present study. This website provides fast queries about the rank of abundance for any gene or protein of each cell line with the whole genome-wide abundance distribution as background, a gene-set based heatmap illustrating the relative abundance of gene expressions between HeLa cells tested, and the correlation scatterplot of HeLa cells displaying the values of any two of the four omics layers. Moreover, all the raw abundance tables at each layer for any gene or genes queried can be easily downloaded using gene symbol or SwissProt ID (**Supplementary Figure 13**). We believe that this website, together with the knowledge which can be derived from it (**Supplementary Figure 14**) will provide beneficial for future research into specific genes and their behavior in complex systems.

## Discussion

Reproducibility is a cornerstone of scientific research and the reproducibility of research results in the life sciences has been questioned. Irreproducible research results are frequently unjustifiably interpreted as the results of incompetence or fraud. In an attempt to bring the discussion to a rationale, evidence-based level the overall problem has been broken down into several specific issues^5–8^. Some of these have been already addressed by the community and have led to changes in research and publication practice. These include the more detailed documentation and validation of research reagents, data collection methods, data openness and transparency, and statistical analyses^7, 73^.

In this paper we study an additional, as yet poorly documented issue affecting reproducibility of research results, the effects of genomic variability in commonly used cells lines on gene expression patterns and phenotypes. Genomic variability is an inherent essential property of evolving biological systems. Therefore, whereas the results of this study were generated from cultured cancer cell lines, they are likely also of significance to the study of populations of single or multicellular organisms, particularly in the context of the emerging personalized/precision paradigm in medicine^74^ The importance of reporting research results in the context of a well defined genotype has recently been highlighted by ageing research in *C. elegans^75^.* Initially a study published in 2010^76^ found as main conclusion that treatment of adult hermaphrodite worms with SOD/catalase mimetics caused a large increase in lifespan. This result could not be reproduced by peers^77^ an situation that was recently explained by the discovery that genetic variation of *C.elegans* strains between labs at least partially explained the previously anecdotal observations of variation in lifespan outcomes^75, 78^.

Like *C.elegans,* human cancer cell lines are another broadly used material in biological and biomedical studies. In most of these studies, cell lines are assumed to be genetically stable rather than heterogeneous at the population level^34^ Our data suggest molecular variability of a cell line as a significant source of apparent irreproducibility even if all other parameters of an experiment are correctly reported and equivalently performed between different labs. The data also show that in HeLa cells, and by inference in other cells with instable genomes, the molecular variability is rapidly evolving, challenging the ability to generate reproducible research results over time even in the same lab. The validation of research reagents and materials is also crucial as it lays the experimental foundation. Previously, discussions around the variability of cells as a source of irreproducible results have centered around experimental error like mislabeling misidentification of cells^12–15^. In this report we focus on the fundamental issue of biological variability which we believe to be more prevalent and significant than simple experimental errors.

Previously genome sequencing studies were reported on HeLa Kyoto^29^ and HeLa CCL2^30^ strains. In this study, we performed a proteome centric, multi-omics investigation of HeLa cells across 13 labs worldwide to evaluate their heterogeneity by focusing on CNV related dosage changes along the axis of gene expression. We find that the HeLa cells tested show significant genomic variability and rapid evolution of genomic variability, likely due to genomic instability. We further show complex, non-linear translation of genotypic variability into expressed transcript and protein patterns. Finally, we document the phenotypic consequences of the proteomic variability among these HeLa cells.

The diverse gene expression patterns among cells named “HeLa” from different labs are not unexpected when one considers the long-term observation of genome instability of HeLa cells^18, 25–28, 79^. In fact, as early as 1979, different strains of HeLa were reported to differ substantially in sensitivity to monofunctional alkylating agents, and the speculated reason was mutation induction in a line of HeLa cells^80^. However, there are no investigations to assess the molecular nature of such differences at a systems-wide scale. Indeed, although HeLa cells are widely used and yield ~0.3% of the total publications listed in PubMed, only a tiny fraction of these papers (< 2% based on keyword search in PubMed) clearly indicated whether they used HeLa Kyoto or HeLa S3. For most of the 97,0 publications, researchers simply report “HeLa” in their method section.

Our study extends the molecular analyses to the assessment of genotypic, proteotypic and phenotypic diversity and has specific implications for how the HeLa cell identities should be documented in future scientific publications. *Firstly,* we found that HeLa CCL2 and Kyoto cells are consistently different at all layers of biology. There are significant differences in cell morphology, doubling time, karyotypes, steady state mRNA, protein expressions, and protein turnover rates. Furthermore, the difference between HeLa CCL2 and Kyoto also results in distinct proteomic responses to *Let7* transfection which can tune protein synthesis from thousands of genes^81^. The gene expression variance between CCL2 and Kyoto is as large as that between many other cell lines of different tissue origins (**Figure 2m**). Therefore, we strongly suggest that all future HeLa related studies should at least clearly report the identities of CCL2 or Kyoto for the cells used. *Secondly,* HeLa S3, the third lineage of HeLa CCL2, did not show distinguishable pattern from CCL2 lines in most of our measurements, which might suggest that HeLa S3, when used, could achieve higher reproducibility to HeLa CCL2 than Kyoto cells. *Thirdly,* we discovered substantial, biologically meaningful divergence of HeLa CCL2 cells after 3 months of culturing that involved copy deletions or gains of some whole chromosomes, 6-7% of gene differential expressions, and quantitative changes of pathways. This suggests that even during studies of moderate duration the molecular makeup of the cells will change. Hence, we recommend the use of early passages of HeLa CCL2 cells directly from the providers like ATCC or minimally of cells with well-documented history of culturing. *Finally,* phenotypes such as growth rate and significant resistance to *S.Tm* infection could be linked to proteotypes (**Figure 5**). Thus, it seems conceivable to establish a genome-wide benchmark to a standard HeLa cell line before performing critical experiments. We herein suggest fast proteome profiling and the quantitative comparison to our results along with the present publication would be a cost effective means to document the biochemical state of the specific cells reported in a publication.

We further suggest that the findings described here derived from HeLa lines are likely generally applicable to other cell lines with instable genomes. For example, the human breast cancer cell line MCF-7, initially derived in 1973 from a malignant pleural effusion, is also a heterogeneous line. As early as 1987 the striking difference of chromosome analysis and the cloning efficiency of four MCF-7 cells from different labs was documented^82^. Compelling ploidy and signaling pathway evidence have been shown for the existence of the triple-negative variants in the MCF cell population^83^. The MCF-7 cells selected for tamoxifen resistance were reported to acquire new phenotypes differing in DNA content, phospho-HER2 and PAX2 expression, and rapamycin sensitivity^84^ Another cell example is HEK293, for which the dynamics of aneuploidy genome in response to cell biology manipulations was discovered by whole-genome sequencing^85^. It is thus imperative to revisit the cell identity and heterogeneity for common cell lines with significant genome instability^79^.

Compared to the genotype, the proteotype provides a close, real-time, and functional snapshot for the better understanding of phenotypes in cancer cells or clinical tissues^37^ Protein level evidences are widely used by antibody-based technologies such as Western blotting in the enormous literatures, and therefore easily communicative. At present, compared to genomics and transcriptomics profiling, large-scale protein level measurements (i.e., proteomics) have been less frequently reported, also due to technical complexities and challenges. SWATH-MS is a new data independent acquisition technique which combines the comprehensiveness of traditional shotgun proteomics, and the reproducibility and quantification accuracy of the targeted proteomics based on SRM or PRM^38, 86,87^. Besides the high proteome coverage of ~5000 proteins quantified, SWATH-MS shows reproducibility performance that is as high as that of transcriptomics to clearly distinguish whole process replicates from different HeLa strains between labs (**Figure 2c**). The harnessing of SWATH-MS, pyProphet FDR control^41^, pSILAC, and other strategies essentially broke the technology hurdle to propel the successful evaluation of steady state proteome expression, protein turnover, and responsive proteome diversities between 14 HeLa cells across labs.

Finally, the multi-omics data for HeLa cells generated in this study provide a high quality resource for understanding gene expression and interplay between layers exemplified by the HeLa Proteome. The HeLa cells from different labs essentially present a cell line panel in which the gene dosages are significantly different whereas the sequence variances are minimal or modest between cells. The total of thousands of dosage changing events observed at each layer maximized the possibility of revealing gene dosage effects during gene expression, and strongly supports the protein complex stoichiometry control as the post-translational regulation contributing to the protein buffering. Also, proteome dynamics of different cellular compartments appeared linked to biological diversity between HeLa cells. However, it remains to be explored whether these regulatory patterns are conserved among other different cell lines.

In conclusion, we comprehensively assayed the genome, transcriptome, and proteome dosages, as well as proteostasis and responsive changes for 14 HeLa cell variants across labs, and demonstrated that the HeLa cells from different providers can yield diverse gene expressions and varied biological experiment results even under identical culturing conditions, and thus provided a new angle for understanding the broad reproducibility crisis.

## Method

### Collection of cells

Frozen HeLa cells were prepared in each laboratory and sent to the coordinating lab at ETH Zurich in dry ice for centralized culturing. A uniform protocol was used at each site to prepare the cells for shipment. Cells collected from a 15 cm dish were transferred into 1 mL freezing media (70% volume of DMEM; 20% FBS; 10% DMSO). Cells were then placed in −20 °C for 2-3 hours, transferred to - 80 °C for 24 hours, and stored in liquid nitrogen until delivery. The frozen state of the delivered cells and existence of dry ice were confirmed upon arrival. Additionally, two aliquots of cell pellets were prepared at each site. The HeLa strains were then centrally cultured in the coordinating laboratory using DMEM medium using standard culturing method according to ATCC.

### Determination of Cell Doubling Time

The cell doubling time for each HeLa strain was determined using a cell counting CCK-8 Kit (Dojindo Laboratories, Japan). Cells were seeded in triplicate at a density of 2800 cells per well of a 96-well plate, and samples were prepared for counting at 5 different time points, 2h, 11h, 24h, 48h, and 72h, respectively. The final doubling time was then calculated (using http://www.doubling-time.com/compute.php). The entire experiments were repeated on two cell lines. Cell doubling time differences between whole-process replicates were all less than 2 hours.

### Phenotypic characterization of HeLa cells

The plates of HeLa cells where imaged with Molecular Devices ImageXpress microscopes, using MetaXpress software. Imaging settings where adjusted for the highest exposure that did not incur over-exposure, with 14 bit dynamic range and laser-based focusing. We imaged 9 sites per well in a 3×3 grid with no spacing and no overlap. Three channels where imaged, with filters for DAPI for imaging the nucleus, GFP for bacteria, and RFP for F-actin. Robotic plate handling was used to load and unload plates (Thermo Scientific).

### Array-CGH (aCGH) analysis

Array-CGH analyses were performed using the Agilent Human Genome CGH Microarray Kit G3 180K (Agilent Technologies, Palo Alto, USA) with 13 Kb overall median probe spacing (Control is DNA pool of 7 normal diploid individuals). Labeling and hybridization were performed following the protocols provided by the manufacturer.

### aCGH processing and gene copy number detection

aCGH measurements of 173,540 genomic probes were used to compute an aCGH log_2_-ratio profile that compares the DNA copy number of each probe in a specific HeLa cell line to normal diploid DNA. aCGH profiles were sorted according to the chromosomal locations of probes and further segmented into chromosomal regions of constant copy number using DNAcopy^88^ (R package DNAcopy with settings smooth.region = 3, outlier.SD.scale = 0.5, smooth.SD.scale = 0.25). Copy number values of individual genes (30,237 known canonical genes of hg19/GRCh37) were determined by mapping chromosomal location of genes to obtained aCGH segments. If a gene was covered by a whole segment, then its copy number value was set to the segment-specific log_2_-ratio. If a gene was covered by more than one adjacent segments (break points within a gene), then its copy number value was set to a weighted average of log_2_-ratio of involved segments according to their overlap with the gene. Most parts of the HeLa genomes are known to be triploid^29, 30^. This was also reflected in our aCGH profiles, but the position of the closest peak to the triploid state varied among the different HeLa cell lines. Therefore, we further aligned the obtained gene copy number measurements to ensure that this peak was located at the triploid state for each cell line.

### RNA extraction, library preparation and mRNA sequencing

Total RNA was collected using the TRIzol^®^ reagent (Life Technologies) following the manufacturer’s instructions. RNA quality was verified on the Agilent 2100 Bioanalyzer (Agilent) and quantity was measured on a Qubit^®^ instrument (Life Technologies).

Libraries were prepared with 4 μg of total RNA using TruSeq RNA kit (Illumina) according to manufacturer’s instructions. Libraries were sequenced on an Illumina HiSeq2000 machine as 100 bp reads single-end. The reads were aligned to the hg19 human genome using TopHat^89^ with standard configurations (no more than 2 mismatches allowed). The numbers of reads for the genes are calculated using the GENCODE v24 release. Only uniquely mapped reads were included.

### Deep analysis of single nucleotide variants and cell line authentication

The RNA sequencing (RNAseq) result of all HeLa cell lines as well as one Hek293 cell line were analyzed for small sequence variants (i.e. single nucleotide variants (SNV) and small insertions/deletions (INDEL)). Raw RNAseq reads for the HEK293 cell line were publicly available and obtained through the NCBI sequence read archive^90^ (accession SRX1300887, downloaded at 2017-01-17 from https://www.ncbi.nlm.nih.gov/sra?term=SRP064410). All RNAseq reads were manually checked with the FastQC^91^ tool (version 0.11.4). Then reads were quality trimmed with Trimmomatic^92^ (version 0.35) and a second quality check was performed with FastQC. Alignment to the *GRCh38* reference genome was performed with STAR^93^ (version 2.4.2a).

Single nucleotide variants were called individually for every cell line, according to a best practices recommendation within the GATK framework^94^ (https://www.broadinstitute.org/gatk/guide/article?id=3891). The pipeline marks duplicate reads in the alignment file with the Picard tools (http://broadinstitute.github.io/picard, version 2.0.1)

MarkDuplicates function. Then the following GATK^94^ (version 3.7), the following tools are sequentially employed: SplitNCigarReads, HaplotypeCaller, VariantFiltration. Variants were filtered according to the following criteria: *Fisher Strand* value above 30, *Quality by Depth* value greater than two and clusters no more than two variants within a 35 base-pair window.

Pairwise cell line concordance was determined as previously described^49^, using in-house R scripts. Briefly, for every cell line pair, concordance marked the fraction of identical variant calls among all variant calls between the cell lines.

### Transfection of Let7d mimics to HeLa cells

Lipofectamine™ RNAiMAX Transfection Reagent (ThermoFisher Scientific) was used to deliver the microRNA human *Let7d* mimics (CAT No. 4464066) as *Let7* treatment, Negative Control (4464058), and the Positive Control of *Kif11* (4390824) provided by Life Technologies Europe (Zug, Switzerland) to all the HeLa strains expect for one GFP positive stain (HeLa 6). The ratio of RNAiMax and DMEM was kept at 1/500 (v/v). The initial seeding concentration was 40 cells /uL for all the strains with the working concentration of miRNAs was 20 nM. The 6-well plate containing 2 mL final transfection medium per well and the 96-well plate containing 100 μL DMEM medium were used for perturbed proteomic measurements and *S.Tm* infection respectively.

### *Salmonella Typhimurium* (S.Tm) infection

HeLa cells in 96-well plate format were cultured for 72 hours before *S.Tm* infection using the InfectX pipeline as described before. Cells were infected for 20 min with S.Tm, incubated for 3 hr and 40 min in medium with 400 μg/ml gentamicin, fixed by 4% PFA, 4% sucrose, and stained with DAPI and DY-547-phalloidin. All liquid handling steps including the infection, fixation, and staining were performed manually. The high-throughput image acquisition was performed using the Molecular Devices ImageXpress microscope (10X S Fluor). It was tested whether let7 mimics would have a phenotype regarding the “docking” step of Salmonella infection. To do so, a non-invasive *S.Tm* strain was used (S.TmΔ4, which lacks the four main SPI-1 effectors SopE, SopE2, SipA and SopB) and added to the cells for 6 min. After that, HeLa cells were 3x washed and the remaining (bound) Salmonella were fixed and then stained to detect them through the automated image analysis. Cells were infected with S.TmΔ4 at an m.o.i. of 125 for 6 min at 37°C and 5% CO2. This non-invasive mutant strain allows to measure the binding capacity of S.Tm. Afterwards, the cells were washed 3 times with 60 μl DMEM/10%FCS and fixed with 60 μl 4% PFA. To visualize bound S.TmΔ4, immunofluorescence staining was applied by the use of a primary anti-LPS antibody (Difco) and a FITC-coated secondary antibody (Jackson). Afterwards, cells were permeabilized and then nuclei were stained with DAPI. A CellProfiler-based image analysis pipeline was applied to determine the infection rate of *S.Tm* for each cell line of the controls and *Let7d* mimics transfected conditions.

### Pulsed SILAC experiment

For the pSILAC experiment, SILAC DMEM High Glucose medium (GE Healthcare) lacking L-arginine, L-lysine was firstly supplemented with light or heavy isotopically labeled lysine and arginine, 10% dialyzed fetal bovine serum (PAN Biotech), and 1% penicillin/streptomycin mix (Gibco). Specifically, 146 mg/L of heavy L-lysine (^13^C_6_^15^N_2_) and 84 mg/L of arginine (^13^C_6_^15^N_4_) (Chemie Brunschwig AG) and the same amount of corresponding non-labeled amino acids (Sigma-Aldrich)^43^ were supplemented respectively to configure heavy and light SILAC medium. Additionally 400 mg/L L-proline (Sigma-Aldrich) was also added into SILAC medium to prevent the potential arginine-proline conversion. HeLa variants were firstly cultured on 15 cm cell culture dishes in pre-prepared, light SILAC medium and stabilized in culture for three to four days. Upon release of cells by 0.25% trypsin/EDTA, cells are respectively counted by a Neubauer hemocytometer. Subsequently, six 10 cm dishes were prepared for each cell variant with a seeding density of 1.5*10^6^ cells per plate, corresponding to three time points with two replicates each. The cell culture plates were incubated for 14 hours, at 5% CO_2_ 37°C, overnight. Cells were then washed three times by PBS at 37°C. The medium was then replaced by Heavy SILAC (K8R10) medium. Cells were harvested and counted in two biological replicates at four different time points (0h, 1h, 4.5h, and 11h). Two dishes of whole process replicate were prepared at each time. The cell pellets were snap frozen in liquid nitrogen after removal of the PBS and stored at −80°C.

### Protein extraction and in-solution digestion

HeLa cells and cell pellets were harvested from shipped tubes, centrally cultured conditions, *Let7d* treated and control experiments, and pSILAC experiment were suspended in 10M urea lysis buffer and complete protease inhibitor cocktail (Roche), ultrasonically lysed at 4°C for 2 minutes by two rounds using a VialTweeter device (Hielscher-Ultrasound Technology). The mixtures were centrifuged at 18,000 g for 1 hour to remove the insoluble material. The supernatant protein amount was quantified by Bio-Rad protein assay. Protein samples were reduced by 10mM Tris-(2-carboxyethyl)-phosphine (TCEP) for 1 hour at 37°C and 20 mM iodoacetamide (IAA) in the dark for 45 minutes at room temperature. All the samples were further diluted by 1:6 (v/v) with 100 mM NH4HCO3 and were digested with sequencing-grade porcine trypsin (Promega) at a protease/protein ratio of 1:25 overnight at 37°C. The 96-well plate format based digestion was used to increase the experimental reproducibility. The amount of the purified peptides was determined using Nanodrop ND-1000 (Thermo Scientific) and 1 μg peptides were injected in each LC-MS run.

### Shotgun proteomics on TripleTOF mass spectrometer

The peptides digested from cell lysate derived from the first time point samples in pSILAC experiment were measured on an SCIEX 5600 TripleTOF mass spectrometer operated in DDA mode ^38, 39,95^, the mass spectrometer was interfaced with an Eksigent NanoLC Ultra 1Dplus HPLC system. Peptides were directly injected onto a 20-cm PicoFrit emitter (New Objective, self-packed to 20 cm with Magic C18 AQ 3-μm 200-Å material), and then separated using a 90-min gradient from 5-35% (buffer A 0.1% (v/v) formic acid, 2% (v/v) acetonitrile, buffer B 0.1% (v/v) formic acid, 100% (v/v) acetonitrile) at a flow rate of 300 nL/min. MS1 spectra were collected in the range 360-1,460 m/z with 250ms per scan. The 20 most intense precursors with charge state 2-5 which exceeded 250 counts per second were selected for fragmentation, and MS2 spectra were collected in the range 50–2,0 m/z for 100 ms. The precursor ions were dynamically excluded from reselection for 20 s.

### SWATH mass spectrometry

Normal proteome, *Let7d* treated and controls, as well as pSILAC samples were all measured by SWATH-MS. The same LC-MS/MS systems used for DDA measurements on SCIEX 5600 TripleTOF was also used for SWATH analysis^38, 95,96^. For normal proteome samples and pSILAC samples a 90-min LC gradient was used, whereas a 60-min gradient were used for the two biological replicates of Let7d experiment. Specifically, in the present SWATH-MS mode, the SCIEX 5600+ TripleTOF instrument was specifically tuned to optimize the quadrupole settings for the selection of 64 variable wide precursor ion selection windows. The 64-variable window schema was optimized based on a normal human cell lysate sample, covering the precursor mass range of 400–1,200 m/z. The effective isolation windows can be considered as being 399.5~408.2, 407.2~415.8, 414.8~422.7, 421.7~429.7, 428.7~437.3, 436.3~444.8, 443.8~451.7, 450.7~458.7, 457.7~466.7, 465.7~473.4, 472.4~478.3, 477.3~485.4, 484.4~491.2, 490.2~497.7, 496.7~504.3, 503.3~511.2, 510.2~518.2, 517.2~525.3, 524.3~533.3, 532.3~540.3, 539.3~546.8, 545.8~554.5, 553.5~561.8, 560.8~568.3, 567.3~575.7, 574.7~582.3, 581.3~588.8, 587.8~595.8, 594.8~601.8, 600.8~608.9, 607.9~616.9, 615.9~624.8, 623.8~632.2, 631.2~640.8, 639.8~647.9, 646.9~654.8, 653.8~661.5, 660.5~670.3, 669.3~678.8, 677.8~687.8, 686.8~696.9, 695.9~706.9, 705.9~715.9, 714.9~726.2, 725.2~737.4, 736.4~746.6, 745.6~757.5, 756.5~767.9, 766.9~779.5, 778.5~792.9, 791.9~807, 806~820, 819~834.2, 833.2~849.4, 848.4~866, 865~884.4, 883.4~899.9, 898.9~919, 918~942.1, 941.1~971.6, 970.6~1006, 1005~1053, 1052~1110.6, 1109.6~1200.5 (containing 1 m/z for the window overlap). SWATH MS2 spectra were collected from 50 to 2,000 m/z. The collision energy (CE) was optimized for each window according to the calculation for a charge 2+ ion centered upon the window with a spread of 15 eV. An accumulation time (dwell time) of 50 ms was used for all fragment-ion scans in high-sensitivity mode and for each SWATH-MS cycle a survey scan in high-resolution mode was also acquired for 250 ms, resulting in a duty cycle of ~3.45 s. A 100 fmol beta-galactosidase standard digest (SCIEX) was injected in-between each run to monitor the instrument performance and to tune the mass accuracy of MS1 and MS2 signals in a real-time manner throughout the sample acquisition.

### SWATH-MS data extraction of protein expression data

With the exception of pSILAC experiment derived data set, all the other SWATH-MS data sets were analyzed and identified by OpenSWATH software^47^ searching against a previously established SWATH assay library which contains mass spectrometric assays for 10,000 human proteins^46^. Profile-mode .wiff files from shotgun data acquisition were centroided and converted to mzML format using the AB Sciex Data Converter v.1.3 and converted to mzXML format using MSConvert v.3.04.238 before OpenSWATH analysis. OpenSWATH firstly identified the peak groups from all individual SWATH maps with statistical control (see below) and then aligned between SWATH maps using a novel TRIC (TRansfer of Identification Confidence)^48^. For large-scale targeted proteomics, protein FDR control needs specific attention and should be equally important compared to shotgun proteomics^41, 46^. Therefore, to pursue a strict statistical quality control of peptide and protein-level identification, we used the newly developed PyProphet extended version^41^ in the present study. This new version of PyProphet combines the set of scores from OpenSWATH for each peptide query to a single discriminant score by applying semi-supervised learning to best separate decoys from high-scoring targets. Particularly for all the label-free samples, PyProphet was run for conducting g-value estimation^41^ for all runs (run-specific context) and in a global fashion (global context). A strategy of two steps of filtering was used: 1) For proteins accepted by PyProphet at FDR <1% in the global context, the sets of peak groups detected at 1% FDR in the run-specific context were included for quantification – this criteria yielded 4335 proteins; 2) For proteins accepted with an FDR <5% in the global context, only those peak groups detected at 1% FDR and also identified in >25% of the total MS runs were accepted. The requantification feature in OpenSWATH was enabled for the filtered protein list. Totally 50,225 peak groups were identified, corresponding to 46,951 unique peptides (43, 521 tryptic peptides) assigned to 5,030 unique SwissProt proteins.

To quantify the protein abundance levels across samples, we summed up the most abundant peptides for each protein (i.e., top 3 peptide groups based on intensity were used for those proteins identified with more than three proteotypic peptide signals whereas all the peptides were summarized for other proteins). This allows for reliably estimating global protein level changes as shown in previous studies^39, 97,98^. The protein expression data matrix was log_10_ transformed and quantile normalized for statistical and bioinformatics analysis.

### SWATH-MS data extraction of pSILAC data

The centroid converted mzML files from shotgun analysis of the first time point samples in pSILAC experiment were searched against different engines including using the iPortal platform^99^ to establish the sample specific library for pSILAC data. Published nine HeLa runs included in the PanHuman Library^46^ were also used to generate this library. iPortal utilized iProphet schema^100^ to integrate the search results from X!Tandem, Omessa, Myrimatch, and Comet at a peptide level FDR =1% by target-decoy strategy. Xinteract option was “-dDECOY_ -OAPdlIw”. Specially, peptide tolerances at MS and MS/MS level were set to be 50 ppm and 0.1 Da respectively. Up to two-missed trypsin cleavages were allowed. Oxidation at methionine was set as variable modification whereas carbamidomethylation at cysteine was set as fixed modification. The iPortal identification result finally contained 3,973 proteins and 42,236 peptides at 1% protein FDR cutoff.

The light version of consensus spectral library was generated using SpectraST^101^. Then the spectrast2tsv.py function in OpenSWATH^47^ was used to generated the light and heavy MS assays as the final library (including the decoy transtions) constructed from top 6 most intense y-ion fragments with Q3 range from 400 to 1200 m/z excluding those falling in the precursor SWATH window were used for targeted data analysis of SWATH maps. OpenSWATH analysis was run with target FDR of 1% and extension FDR of 5%^47^ (quality cutoff to still consider a feature for alignment) and aligned by TRIC^48^, whereas requantificated data points were discarded for protein turnover calculation.

### Determining protein turnover rates

In pSILAC, the quantification of heavy and light signals of proteins at different timepoints after pulse labeling permits the quantification of protein specific turnover rates. The protein turnover rate was determined similarly to our previous study^45^. The rates of loss of the light isotope (k_loss_) were directly calculated from the output data matrix generated by OpenSWATH, following the methods previously described by Pratt et al^102^: specifically, we modeled the Relative Isotope Abundance (RIA), defined as the signal intensity in the light channel divided by the sum of light and heavy intensities, onto an exponential decay model assuming a null heavy intensity (RIA = 1) at time 0, i.e., RIA(t)= *e*^*(~kloss×t)*^

An in-house package was used to perform the fit that was obtained through nonlinear least-squares estimation^45^ and to further perform a weighted average of the k_loss_ value of each peptides of the protein, which ensured giving more weight to peptides carrying robust information. Importantly, this method maximized correlation and minimized median absolute error across replicates as described before^45^.

Boisvert et al^103^ reported a significant recycling of the unlabeled amino acid from the light protein degradation in HeLa cells where they had to use a 0.8 offset to calculate the protein degradation rate (Kdeg) from Kloss. Therefore in the present study we simply used the direct “proxy of turnover rate” i.e., *log2* (*K_loss_*), whenever applicable, to perform the cross-cell and multi-omics comparisons and illustrations^58^. This proxy parameter essentially avoids the data distortion due to the possible different light amino acid recycling speed and inaccuracy of cell doubling time determination. Only those Kloss values assayed in every cell sample were accepted for cross-comparison.

### Estimation of absolute protein copies

HeLa cells are a heavily investigated cell line for which many research resources are available. In a study published by Zeiler et al^104^, the absolute protein copy numbers in a HeLa cell line (unknown identity) were determined using Protein Epitope Signature Tag (PrEST) and SILAC-based absolute quantification^104^. For all the anchor PrEST proteins in Zeiler et al^104^ we covered the full dynamic range in this study. This enabled us to correlate 23 HeLa anchor proteins with their respective copy numbers reported in Zeiler et al to our summed top three peptide SWATH-MS intensity estimates for each protein. The high Pearson correlation coefficient (average R=0.872) allowed us to directly utilize the correlation equation in each cell line to infer the protein copy numbers for all the protein identified by OpenSWATH.

### Other bioinformatic analyses

To calculate the Kyoto/CCL2 ratios at all levels, HeLa 11 was excluded due to its deviating genome dosage type (**Figure 1B**), so that there are six cell variants in each group evenly. Cellular compartment annotated was done by mapping protein identifies to a recently established subcellular map of human proteome^61^. The David Bioinformatics v6.8 (https://david.ncifcrf.gov) was used to exact the protein annotations for other organelles of interest that is not covered by subcellular atlas^61^. Protein complexes information were directed extracted from CORUM database^105^ and mapped through SwissProt ID. The t-SNE plots are done using the R package Rtsne version 0.13. As parameters, we used a perplexity of 20 and 500 iterations. PCA are done using the R package FactoMineR version 1.39 (We chose 5 principal components). For differential gene expression analysis between HeLa 11 and other cells, we fitted a linear model using the R package limma version 3.30.13 then we corrected by false discovery rate with adjusted p value at 0.05. The gene set enrichment analysis was performed by STRING database^106^ (https://string-db.org). For the circle plots, we used R package RCircos version 1.2.0 and initialized the cytoband with UCSC.HG19.Human.CytoBandIdeogram. Heatmaps are made using R package pheatmap version 1.0.8. For the website, we used the R packages shiny version 1.0.5, shinydashboard version 0.6.1 and shinyBS.

## Author Contributions

Y.L. and R.A. designed and over saw the project. Y.L., Y.M., E.G.W., P.L.G., M.F., I.B., M.S., M.E., and F.B. analyzed the data performed the bioinformatics analysis. Y.M. developed the HeLa Proteome Web. T.M. performed pulse SILAC experiment. S.K. and Y.L. performed Let7 experiment. S.K. performed S.Tm infection experiment. A.V.D., C.B, I.S., C.D., and H.Z. established and cultured the cell lines. Y.L. and M.M performed the mass spectrometry experiments. I. B. performed pyProphet analysis. F.S.B. generated CNV data. M.S. processed the CNV data. C.B. generated RNA-seq data. M.F. performed sequence variation analysis. F.B. and P.L.G analyzed RNA-Seq data. M.E. analyzed the microscopy phenotypic data. G.T. and J.S.R supervised data interpretation. S.E.A. supervised the genomics data generation. W.D.H. supervised all the microbiology experiments and provide critical inputs. Y.M., T.M., S.K. had comparable contribution to the paper. Y.L., E.W.G., and R.A. wrote the paper.

## Acknowledgement

We thank Andreas Beyer, Ben Collins, Sergey Nikolaev, George Rosenberger for helpful discussions. We thank Hui Zhang and Jing Chen from Johns Hopkins University, Delphine Pflieger and Odile Filhol-cochet from CEA Grenoble, Meliana Riwanto from University Hospital Zurich, Urs Greber and Maarit Suomalainen from University of Zurich, Cécile Arrieumerlou from University of Basel (through InfectX), Martin Beck and Marie-Therese Mackmull from European Molecular Biology Laboratory, Claus Jorgensen and Jonathan Worboys from Cancer Research UK Manchester Institute, Matthias Peter and Chris Barnes from ETH Zurich, Ashok Venkitaraman and Cigdem Williams from University of Cambridge for providing us the HeLa cells they used.

The work was supported by the SystemsX.ch project PhosphoNetX PPM (to R.A.) and TargetInfectX (to C.D.), the Swiss National Science Foundation (grant no. 3100A0-688 107679 to R.A.), the European Research Council ERC-20140AdG 670821 to R.A.), the JRC for Computational Biomedicine which was partially funded by Bayer AG to J.S.R., Swiss National Science Foundation (grant no. 163180 to S.E.A.), the European Research Council AdG 249968 to S.E.A.), the European Research Council (ERC grant number 616441-DISEASEAVATARS to G.T.), the Umberto Veronesi Foundation (fellowship to P.L.G.), the ERA-NET Neuron Program (P.-L.G.), Regione Lombardia (Ricerca Indipendente 2012 to G.T.), and Italian Ministry of Health (Ricerca Corrente to G.T.) Evan Williams was supported by an NIH F32 Ruth Kirchstein Fellowship (F32GM119190). Yansheng Liu was awarded by the start-up funding for Assistant Professor from Yale Cancer Biology Institute and Yale University School of Medicine.

